# Comparable analysis of multiple DNA double-strand break repair pathways in CRISPR-mediated endogenous tagging

**DOI:** 10.1101/2023.06.28.546861

**Authors:** Chiharu Tei, Shoji Hata, Akira Mabuchi, Shotaro Okuda, Kei K Ito, Mariya Genova, Masamitsu Fukuyama, Shohei Yamamoto, Takumi Chinen, Atsushi Toyoda, Daiju Kitagawa

**Affiliations:** Department of Physiological Chemistry, Graduate School of Pharmaceutical Sciences, The University of Tokyo, Bunkyo, Tokyo, Japan; Precursory Research for Embryonic Science and Technology (PRESTO) Program, Japan Science and Technology Agency, Honcho Kawaguchi, Saitama, Japan; Zentrum für Molekulare Biologie, Universität Heidelberg, DKFZ-ZMBH Allianz, Heidelberg, Germany; Comparative Genomics Laboratory and Advanced Genomics Center, National Institute of Genetics, Mishima, Shizuoka, Japan

## Abstract

CRISPR-mediated endogenous tagging, utilizing the homology-directed repair (HDR) of DNA double-strand breaks (DSBs) with exogenously incorporated donor DNA, is a powerful tool in biological research. Inhibition of the non-homologous end joining (NHEJ) pathway has been proposed as a promising strategy for improving the low efficiency of accurate knock-in via the HDR pathway. However, the influence of alternative DSB repair pathways on gene knock-in remains to be fully explored. In this study, our long-read amplicon sequencing analysis reveals various patterns of imprecise repair in CRISPR/Cas-mediated knock-in, even under conditions where NHEJ is inhibited. Suppression of the microhomology-mediated end joining (MMEJ) or the single strand annealing (SSA) repair mechanisms leads to a reduction in distinct patterns of imprecise repair, thereby elevating the efficiency of accurate knock-in. Furthermore, a novel reporter system shows that the SSA pathway contributes to a specific pattern of imprecise repair, known as asymmetric HDR. Collectively, our study uncovers the involvement of multiple DSB repair pathways in CRISPR/Cas-mediated gene knock-in and proposes alternative approaches to enhance the efficiency of precise gene knock-in.

## Introduction

The CRISPR/Cas-mediated endogenous gene tagging system is an important tool for analyzing protein localization and function in an endogenous context by introducing a DNA sequence encoding for a peptide or a protein tag into the gene of interest. In this system, Cas endonucleases are targeted to the specific gene locus with a guide RNA to induce double-strand DNA breaks (DSBs). Cpf1 and Cas9, which are the most commonly used Cas nucleases, recognize different PAM sequences and cleave DNA in distinct manners (Gasiunas et al., 2012; Jinek et al., 2012; Zetsche et al., 2015). These DSB sites can be repaired through the homology-directed repair (HDR) pathway, which uses a homologous DNA sequence as a template to seal the DSB precisely. Exogenously introduced DNA, containing so-called homology arms (HAs) − elements that are homologous to the region flanking the DSB site, can also serve as a repair template. Therefore, a desired sequence flanked by HAs in such a donor DNA template can be accurately integrated into the specific gene locations via the HDR pathway in the CRISPR/Cas-mediated gene knock-in (Yeh et al., 2019).

Despite the broad applications of this method, the low efficiency of accurate editing events remains a significant challenge (Javaid and Choi, 2021). This issue arises from various DSB repair patterns other than the precise incorporation of the donor DNA sequences, which is referred to as “perfect HDR” (Canaj et al., 2019). For example, DSBs can be repaired without utilizing donor DNA, leading either to a seamless reconstitution of the wild-type (WT) sequence or introducing insertions/deletions (indels) at the lesion locus. Moreover, even in cases where the DSB repair is dependent on the donor DNA, there is a frequent incidence of imprecise donor integration into the target site. The imprecise incorporation can result in various faulty repair patterns, such as partial integration of the donor sequence, HAs duplication, and “asymmetric HDR”, where only one side of donor DNA is precisely integrated but the other is not (Canaj et al., 2019). The non-homologous end joining (NHEJ), which is recognized as the most dominant DSB repair pathway, ligates the free DNA ends at the lesion in a homology-independent manner at the cost of higher indel incidence (Chang et al., 2017). Suppressing this pathway leads to a reduction of indel occurrence, thereby enhancing the efficiency of perfect HDR (Fu et al., 2021; Maruyama et al., 2015). Based on this, a strategy of inhibiting non-HDR repair pathways has been proposed as an effective approach to increase the efficiency of precise knock-in events (Rozov et al., 2019).

In addition to NHEJ and HDR, there are two alternative non-HDR DSB repair pathways: microhomology-mediated end joining (MMEJ), and single strand annealing (SSA) (Ceccaldi et al., 2016; Scully et al., 2019). MMEJ relies on the annealing of two microhomologous sequences (2-20 nt) flanking the broken junction, which frequently results in introduction of deletions at the junction (Sfeir and Symington, 2015; Sinha et al., 2016). While the MMEJ pathway has been regarded as a minor DSB repair pathway compared to NHEJ, there is growing evidence that suppression of the MMEJ pathway by inhibiting its central effector POLQ increases HDR frequency in CRISPR-mediated knock-in (Arai and Nakao, 2021; Schimmel et al., 2023). In contrast, the last DSB repair pathway, the SSA pathway, utilizes Rad52-dependent annealing of longer homologous sequences for repair of DSBs (Bhargava et al., 2016). In the context of CRISPR-mediated gene editing, it has been reported that SSA-mediated repair occurs in artificial gene cassettes containing two homologous regions, resulting in deletions of the intervening sequence between them (Li et al., 2018; van de Kooij et al., 2022). Although these two non-HDR pathways have been suggested to take part in repairing DSBs introduced by Cas nucleases as mentioned, it is not yet clear how they impact the repair outcomes in CRISPR/Cas-mediated knock-in.

In this study, we investigated the contributions of these non-HDR pathways to the repair outcomes of CRISPR/Cas-mediated endogenous gene tagging with fluorescent proteins in human non-transformed diploid cells. Our observation revealed the presence of various imprecise repair patterns along with perfect HDR, even when NHEJ is inhibited. The inhibition of POLQ and Rad52, key components of the MMEJ and SSA pathways, respectively, resulted in a decrease in deletions around the target site, while SSA suppression exhibited a sequence-dependent and specific effect. Furthermore, we found that suppression of the SSA pathway resulted in decreased occurrence of various donor mis-integrations, especially of asymmetric HDR events. Taken together, our findings indicate that the MMEJ and SSA pathways contribute to distinct patterns of imprecise editing, and thus their inhibition can further enhance the efficiency of precise gene knock-in in combination with NHEJ suppression.

## Results

### NHEJ inhibition is not sufficient to completely suppress non-HDR repairs in Cpf1- and Cas9-mediated endogenous tagging

We first examined the effects of the NHEJ inhibition on knock-in outcomes in the hTERT-immortalized RPE1 cell line, which is a human non-transformed diploid cell line commonly used in the field of cell biology. For knock-in in RPE1 cells, we applied a cloning-free endogenous tagging method established previously (Mabuchi et al., 2023). Following this protocol, we prepared donor DNA by PCR using a pair of primers containing 90 bases of HA sequences. Recombinant Cas nucleases and guide RNAs transcribed *in vitro* were mixed to form RNP complexes, which were electroporated into cells along with the donor DNA (Fig. 1a). To examine the effects of the inhibition of NHEJ on both Cpf1- and Cas9-mediated knock-in, we performed a Cpf1-mediated C-terminal tagging of the nuclear protein HNRNPA1 and a Cas9-mediated N-terminal tagging of the trafficking protein RAB11A with the green fluorescent protein mNeonGreen (mNG). Immediately after the electroporation of either Cpf1-RNP or Cas9-RNP along with the donor DNA, these cells were treated with Alt-R HDR Enhancer V2, an NHEJ inhibitor (NHEJi), for 24 hours (Schubert et al., 2021) (Fig. 1a). Fluorescence imaging showed that most of the mNG-positive cells exhibited the expected localization corresponding to each mNG-fused endogenous protein with few exceptions of abnormal mNG signal localization observed in the knock-in at the *RAB11A* locus (Fig. 1b, Fig. S1a). Consistent with previous reports (Fu et al., 2021; Ghetti et al., 2021), NHEJi treatment increased the cell population with mNG signal (Fig. 1b, Fig. S1a). For precise quantification of knock-in efficiency, we performed flow cytometric analysis 4 days after electroporation. This analysis revealed that NHEJi treatment increased knock-in efficiency by nearly 3-fold for Cpf1-mediated knock-in at the *HNRNPA1* locus (3.8% to 11.7%) and nearly 2.5-fold for Cas9-mediated knock-in at the *RAB11A* locus (7.1% to 17.5%) (Fig. 1c, d).

**Fig. 1:**
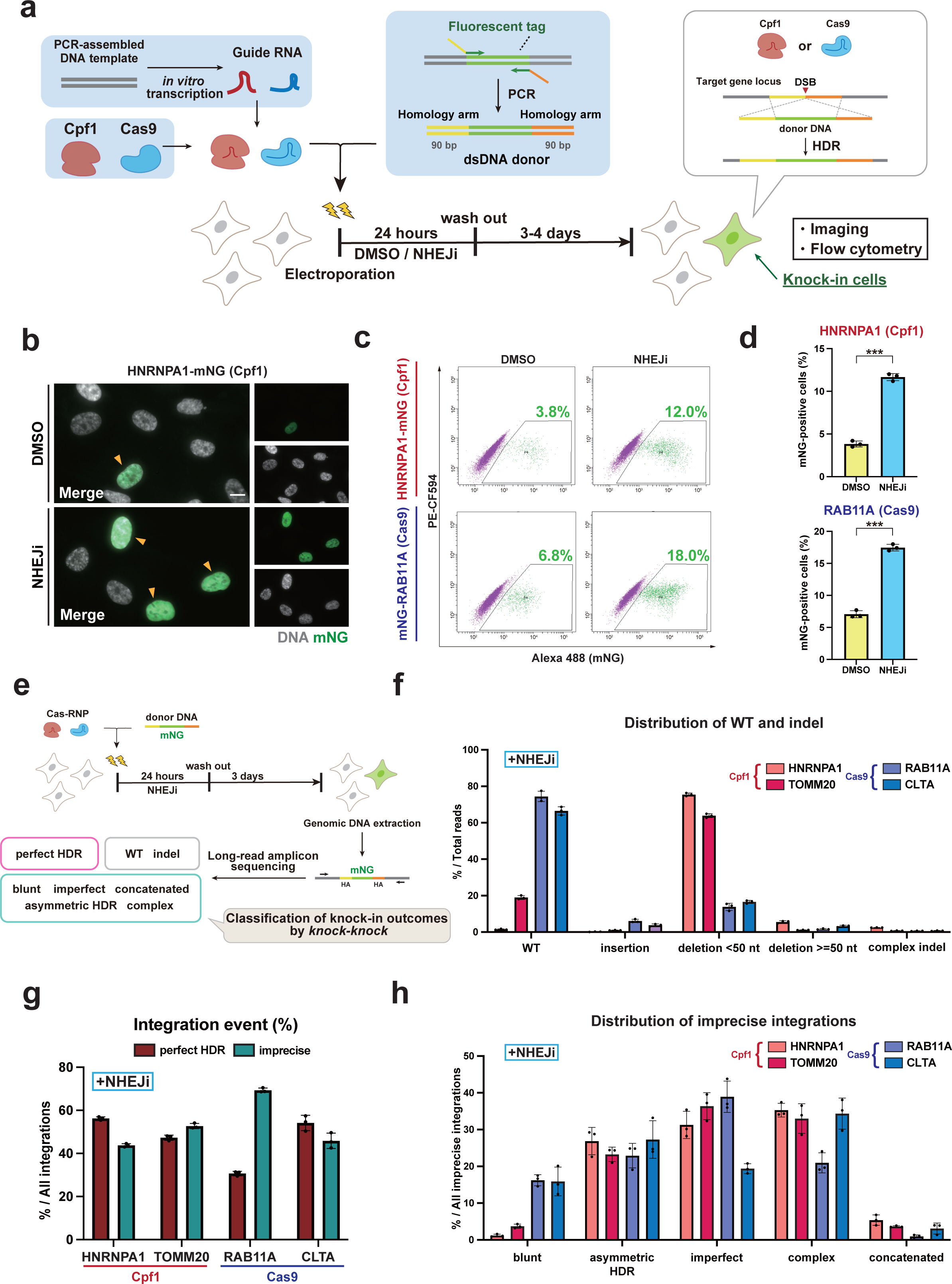
NHEJ inhibition is insufficient for complete suppression of repair outcomes other than perfect HDR. **a**, Schematic overview of endogenous gene tagging in human RPE1 cells. The cells were cultured in a medium containing DMSO or 1 μM NHEJi for 24 hours after electroporation of Cas-RNP and the donor DNA. **b**, Representative images from cells with Cpf1-mediated mNG tagging of HNRNPA1. Arrowheads indicate cells with nuclear mNG signals. Cells at 5 days after electroporation were fixed and analyzed. Scale bar: 10 µm. **c**, Flow cytometric analysis of Cpf1-mediated HNRNPA1-mNG and Cas9-mediated mNG-RAB11A knock-in cells. Cells at 4 days after electroporation were analyzed. Percentages of cells with mNG signal are shown in the plots. **d**, Quantification of percentages of mNG-positive cells from (**c**). Approximately 10,000 cells were analyzed for each sample. **e**, Schematic overview of sequencing-based approach to analyze overall repair patterns of CRISPR-mediated endogenous tagging. **f**, The distribution of repair patterns without donor integration was shown across the targeted gene loci after *knock-knoc*k categorized each sequence read into a specific category of knock-in outcomes and calculated the frequency of each repair category. 122496-211434 reads were analyzed for each sample. **g**, The percentage of perfect HDR and imprecise integration within total integration events for the target genes. For each sample, 2784-30651 reads categorized as the integration events were analyzed. **h**, Distribution of imprecise integration events across the targeted genes. For each category, percentage within total imprecise integration events are shown. In **d**, **f**, **g** and **h**, data from three biological replicates are represented as mean ± S.D. and P-values were calculated by a two-tailed, unpaired Student’s t-test. ***P < 0.001.

Having confirmed that NHEJi has a significant positive effect on the knock-in efficiency, we further checked the resulting repair patterns following the Cas nuclease-induced DNA DSBs at the target loci under NHEJ inhibition. For this purpose, we conducted the long-read amplicon sequencing using PacBio for knock-in alleles followed by genotyping using a computational framework called *knock-knock* (Canaj et al., 2019). We applied this approach to mNG tagging of HNRNPA1 and TOMM20 using Cpf1, and of RAB11A and CLTA using Cas9 under conditions where NHEJ was inhibited (Fig. 1e). After the electroporation followed by NHEJi treatment, the target sites of knock-in were amplified by PCR from extracted genomic DNA. Subsequent to the sequencing process, each Hi-Fi read was categorized into specific DSB repair outcomes, such as WT, indels, perfect HDR, or subtypes of imprecise integration, through the *knock-knock* classification process. Among the cases where the donor DNA was not inserted into the target site, the Cas9-mediated cleavage mostly resulted in WT repair rather than introduction of indels. In contrast, repairs characterized by deletions smaller than 50 nt were the predominant outcome of Cpf1-mediated cleavage (Fig. 1f). Given the comparable knock-in efficiencies for both Cas9 and Cpf1 as shown in Fig. 1d, the differences in the repair patterns can be attributed to the unique DNA DSB ends generated by these two Cas nucleases: blunt ends by Cas9 and staggered ends with Cpf1 (Gasiunas et al., 2012; Jinek et al., 2012; Zetsche et al., 2015).

Intriguingly, the ratio of perfect HDR in all integration events was still far below 100% even upon the inhibition of NHEJ, which is the most dominant non-HDR DSB repair pathway in cells (Fig. 1g). In the knock-in experiments targeting HNRNPA1, TOMM20, and CLTA, about half of all integration events exhibited perfect HDR, whereas for RAB11A, this desired outcome was observed in only 30% of the cases. In all gene loci, there were certain proportions of imprecise integrations. They may result in the expression of the fluorescent protein with the unintended subcellar localization as observed in the mNG tagging of RAB11A (Fig. S1a).

Based on the classification by *knock-knock*, we placed the imprecise integration events into the following categories: blunt (both ends of the donor DNA including the HAs are directly ligated to the cut site), asymmetric HDR (only one side of the donor DNA is precisely integrated in an HDR manner), imperfect (at least one end of the donor DNA is trimmed and incorporated), concatenated (multiple insertions of donors), and complex (not classified into the other four mis-integration categories). Among the imprecise insertions detected in both Cpf1- and Cas9-based endogenous tagging approaches, asymmetric HDR and imperfect integrations were the major patterns, together with complex integrations (Fig. 1h). One difference between the repair patterns of Cpf1- and Cas9-mediated knock-in is that the Cas9-based genome editing tended to have a higher percentage of blunt integration events compared to Cpf1. This observation can also be attributed to the unique DNA DSB ends generated by these two Cas nucleases. Furthermore, even when the same Cas nuclease was used, there were some variations in the repair patterns among the different gene loci. For example, in the Cas9-mediated knock-ins, the frequency of imperfect integration was higher in RAB11A than in CLTA. Conversely, a lower rate of complex integration was observed for RAB11A compared to CLTA. These differences suggest that the repair pattern is dependent on the genomic context of the targeted gene loci in addition to the types of Cas nucleases. Collectively, these data suggest that various imprecise integrations can occur in high frequency via non-HDR pathways other than NHEJ in Cpf1- or Cas9-mediated gene knock-in.

### The MMEJ and SSA non-HDR pathways influence the efficiency of gene knock-in upon NHEJ inhibition

Next, to examine the contribution of minor non-HDR pathways other than NHEJ to the knock-in process, we attempted to suppress the MMEJ and SSA pathways in addition to NHEJ inhibition. For evaluation of bi-allelic knock-in efficiency along with total knock-in efficiency, we performed simultaneous dual-color tagging with both mNG and the red fluorescent protein mScarlet (Fig. 2a). Dual-color tagging of HNRNPA1 resulted in a comparable percentage of total fluorescence-positive cells as single-color tagging of the same gene in both DMSO control and NHEJi conditions (Fig. 2b, c). Interestingly, inhibition of NHEJ had a much higher impact on the percentage of mNG and mScarlet double-positive cells (0.06% to 1.49%, more than 20 times higher) than on that of total fluorescence-positive cells (5.3% to 17.0%, about 3 times higher) in the dual-color tagging (Fig. 2b, c). The marked increase in the frequency of double-positive cells may reflect the non-linear correlation between editing efficiency and the rate of bi-allelic editing, whereby the probability of bi-allelic editing theoretically increases proportional to the square of the editing efficiency (Naert et al., 2020; Takayama et al., 2017). Thus, estimation of bi-allelic knock-in efficiency using the dual-color tagging system is expected to provide a much more sensitive method for evaluating knock-in likelihood compared to the single-color tagging.

**Fig. 2:**
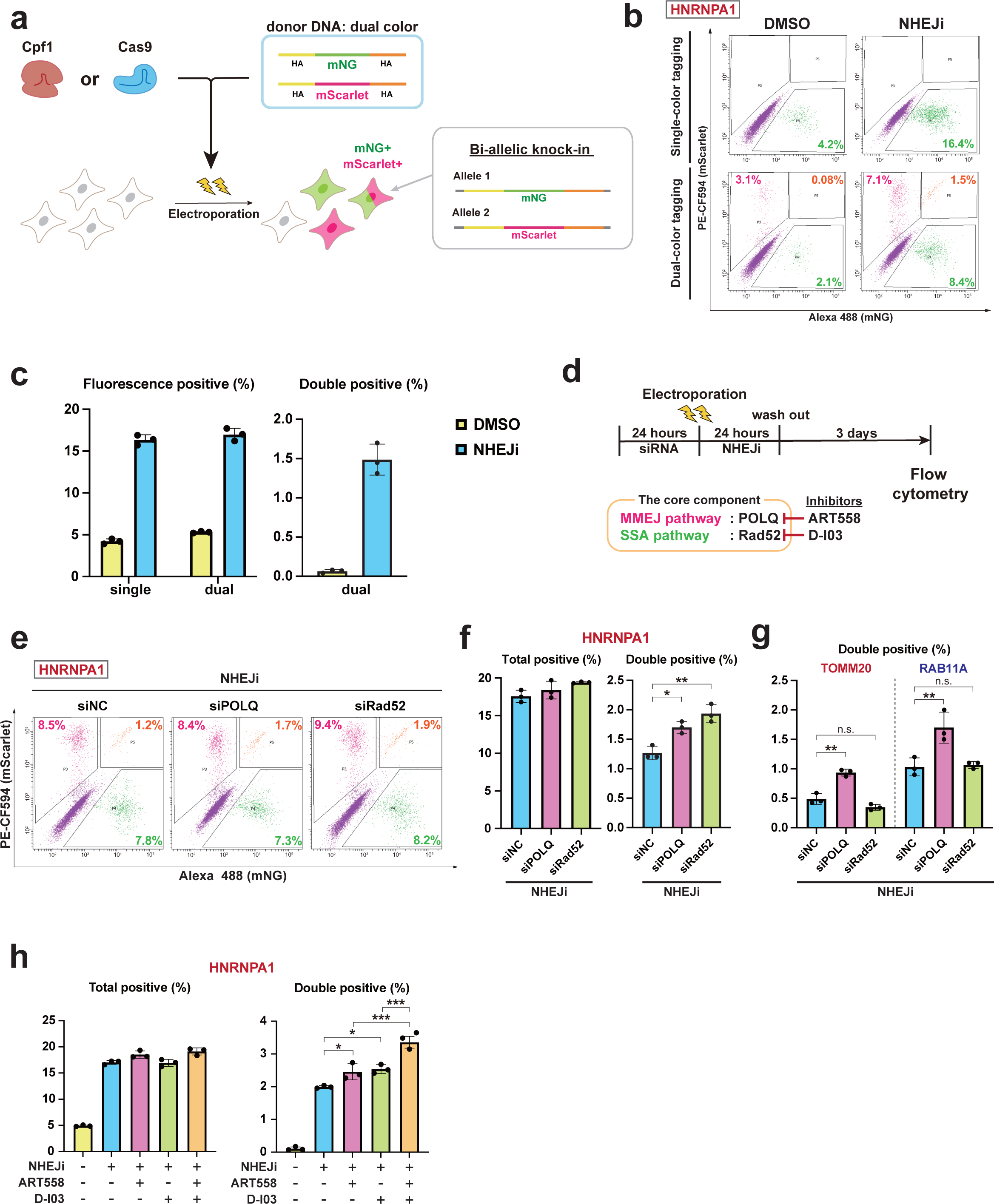
Under the inhibition of NHEJ, the suppression of the MMEJ and SSA pathways affects the efficiency of endogenous gene tagging. **a**, Schematic of dual-color tagging, where two DNA donors containing mNG or mScarlet sequence were simultaneously introduced with CRISPR RNPs. **b**, Flow cytometric analysis for cells with mNG tagging (single-color tagging) and mNG/mScarlet tagging (dual-color tagging) of HNRNPA1. After electroporation, the cells were treated with DMSO or NHEJi for 24 hours and subsequently cultured for additional three days before the analysis. The plots display the percentages of cells exhibiting only mNG signal (green), only mScarlet signal (magenta), or both signals (orange). **c**, Quantification of percentages of fluorescence-positive (left) and double-positive cells (right) from (**b**). In single-color tagging, fluorescence-positive cells refer to mNG-positive cells. As for dual-color tagging, fluorescence-positive cells correspond to those that are positive for either mNG or mScarlet, or both, while double-positive cells refer to those that exhibit positive signals for both mNG and mScarlet. Approximately 10,000 cells were analyzed for each sample. **d,** Schematic diagram showing time course of the dual-color tagging. siRNA transfection was performed 24 hours before electroporation of Cas-RNP and donor DNA. After treatment with 1 µM NHEJi for 24 hours and subsequent incubation in fresh media, cells were subjected to flow cytometric analysis. **e**, Flow cytometric analysis as in (**b**). Cells were treated with the indicated siRNA before electroporation as shown in (**d**). **f**, Quantification of percentages of total- and double-positive cells in dual-color tagging from (**e**). Approximately 10,000 cells were analyzed for each sample. **g.** Quantification of percentages of double-positive cells for the indicated conditions in dual-color tagging of TOMM20 using Cpf1 and RAB11A using Cas9. Approximately 10,000 cells were analyzed for each sample. **h**, Quantification of percentages of total- and double-positive cells in dual-color tagging of HNRNPA1 using Cpf1. Cells were treated with the indicated inhibitors for 24 hours after electroporation. More than 5,000 cells were analyzed for each sample. In **c**, **f**, **g** and **h**, data from three biological replicates are presented as mean ± S.D. and P value was calculated by a Tukey–Kramer test in this figure. *P <0.05, **P <0.01, ***P <0.001, n.s.: Not significant.

For the suppression of the MMEJ pathway, we first used siRNA-mediated gene knockdown of POLQ, which plays the central role in this repair process. Cpf1-mediated dual-color tagging of HNRNPA1 with mNG and mScarlet was performed by electroporation 24 hours after siRNA transfection (Fig. 2d). Cells were subsequently treated with NHEJi for 24 hours and subjected to flow cytometry 4 days after electroporation. Knockdown of POLQ significantly increased the proportion of double-positive cells, while showing a minor effect on the percentage of total fluorescence-positive cells (Fig. 2e, f). In tagging of TOMM20 and RAB11A via Cpf1 and Cas9, respectively, depletion of POLQ also led to an increased rate of double-positive cells (Fig. 2g). The impact of POLQ depletion was further validated by treatment with a recently discovered POLQ inhibitor, ART558 (Zatreanu et al., 2021). Upon ART558 treatment along with NHEJi for 24 hours, the fraction of double-positive cells increased compared to only NHEJi-treated condition at the *HNRNPA1* locus (Fig. 2h). These data indicate that the suppression of the MMEJ pathway has a positive effect on knock-in efficiency.

We next inhibited the SSA pathway by disrupting the function of Rad52, the core component of this repair mechanism responsible for annealing single-stranded DNA (ssDNA) with homologous sequences (Jalan et al., 2019). Under condition of NHEJ inhibition, siRNA-mediated Rad52 depletion significantly increased the population of double-positive cells in dual-color tagging of HNRNPA1 (Fig. 2e, f). Moreover, treatment with D-I03, a specific inhibitor targeting Rad52 (Huang et al., 2016), similarly elicited a notable elevation in the proportion of double-positive cells as for tagging of HNRNPA1 (Fig. 2h). However, the dual-color tagging of the other two loci was not affected by Rad52 depletion (Fig. 2g). These data suggest that the effect of suppressing the SSA pathway on knock-in efficiency varies depending on the target gene. For the *HNRNPA1* locus, which is sensitive to SSA suppression, simultaneous treatment of ART558 and D-I03 synergistically increased the efficiency of the dual-color tagging (Fig. 2h). This observation indicates that the two minor DSB repair pathways contribute differentially to the CRISPR-mediated knock-in process.

### Long-read amplicon sequencing revealed the effects of MMEJ and SSA suppression on repair patterns of Cpf1-mediated endogenous tagging

We next assessed the influence of suppressing the non-HDR pathway on the repair patterns of Cpf1-mediated mNG tagging of HNRNPA1. The long-read amplicon sequencing and subsequent *knock-knock* analysis revealed a noticeable increase in the proportion of perfect HDR within total reads when cells were treated with either the MMEJ inhibitor ART558 or the SSA inhibitor D-I03, under NHEJ inhibition (Fig. 3a). This increase can be attributed to a significant decrease in the proportion of deletions with less than 50 nt observed in both ART558- or D-I03-treated conditions (Fig. 3a).

**Fig. 3:**
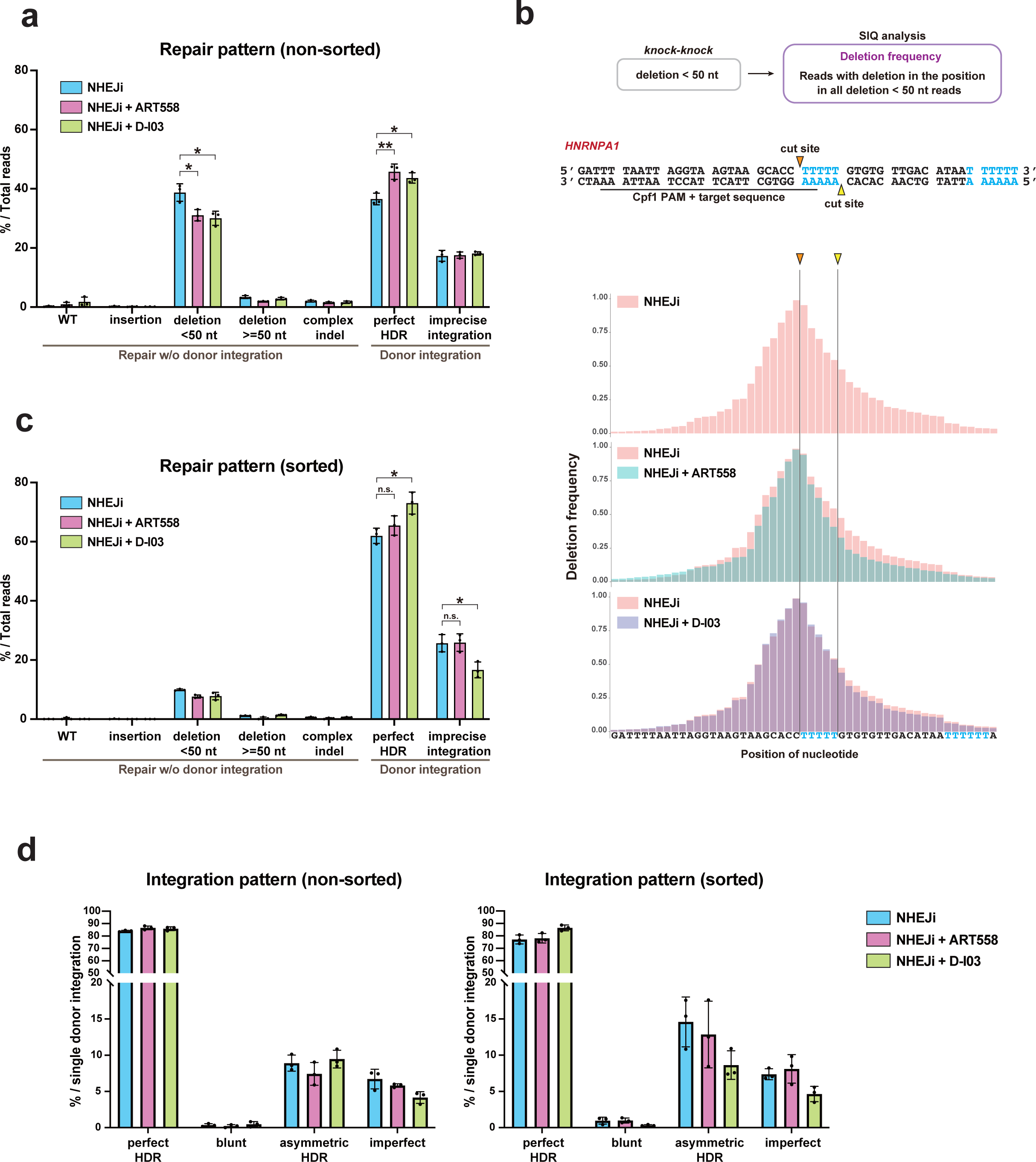
The MMEJ and SSA pathways influence repair patterns of Cpf1-mediated endogenous tagging. **a**, Distribution of repair patterns in Cpf1-mediated mNG tagging of HNRNPA1 in cells treated with the indicated inhibitors for 24 hours after electroporation. Long-read amplicon sequencing and subsequent *knock-knock* analysis were performed following the methodology depicted in Fig. 1e. 115790-230164 sequencing reads were analyzed for each sample. **b**, Positional distribution of deletions within reads categorized as deletions with less than 50 nt by *knock-knock* in (**a**). The percentage of reads with a deletion at each nucleotide was calculated and visualized using SIQ program and SIQPlotteR. A total of 133684-186231 reads from three biological replicates for each condition were analyzed and plotted collectively. **c**, Distribution of repair patterns in mNG tagging of HNRNPA1 within fluorescence-positive cells. Following the same knock-in and incubation procedure as in (**a**), mNG-positive cells were sorted using flow cytometry and subjected to sequencing and *knock-knock* analysis. 147266-226842 reads were analyzed for each sample. **d**, Distribution of donor integration patterns from *knock-knock* analysis of non-sorted (**a**) and sorted cells (**c**). For each category, the percentage within single donor DNA integration events (perfect HDR, blunt integrations, asymmetric HDR, and imperfect integrations) was calculated. Data from three biological replicates are presented as mean ± S.D. and P value was calculated by a Tukey–Kramer test in this figure. *P <0.05, **P <0.01, n.s.: Not significant.

To investigate the specific features of reduced deletions by ART558 and D-I03, we extracted the sequencing reads that knock-knock classified as deletions with less than 50 nt and analyzed them using the Sequence Interrogation and Quantification (SIQ) program, which enabled us to analyze the positional distribution of deletions (van Schendel et al., 2022). Intriguingly, in the condition where only NHEJ was inhibited, the peak of the deletion distribution is biased toward upstream of the cut sites of Cpf1 (Fig. 3b). Since the cut site of Cpf1 in the strand targeted by crRNA is located outside of the target sequence, the DSB can be imprecisely repaired without disrupting target recognition by Cpf1, which is supposed to allow repeated cleavages until the deletion reaches within the target sequence (Zetsche et al., 2015). This unique property of Cpf1 could result in a high frequency of deletions within the target sequence among various other patterns, leading to the biased distribution observed (Fig. 3b) In the context of NHEJ inhibition, ART558 decreased the frequency of deletions spanning a wide range (Fig. 3b). In particular, the deletions in the downstream region of the cut site flanked by two T-rich sequences (indicated in cyan) showed a pronounced decrease. This susceptibility to ART558 suggests that MMEJ-mediated repair becomes prominent around this region, probably facilitated by the availability of multiple combinations for microhomology annealing between the two T-rich sequences. Notably, while ART558-mediated reduction of deletions was observed in other positions as well, D-I03 treatment exhibited a distinct tendency to selectively reduce deletions between the T-rich sequences (Fig 3b). These findings indicate that the suppression of MMEJ and SSA enhances the efficiency of perfect HDR by reducing deletions through distinct mechanisms.

### The suppression of the SSA pathway increases the proportion of perfect HDR events among the fluorescence-positive cells

In the process of establishing endogenously tagged cell lines with fluorescent proteins, flow cytometric cell sorting is a common step to enrich cells with successful knock-in. To examine the effects of MMEJ or SSA pathway suppression in such practical situations, we performed Cpf1-mediated mNG tagging of HNRNPA1 and subsequently collected 10,000 mNG-positive cells using flow cytometry, with which we performed long-read amplicon sequencing and *knock-knock* analysis. The results showed that the overall proportions of the donor-independent repair outcomes, especially deletions of less than 50 nt, significantly decreased compared to those in non-sorted cells (Fig. 3a, c). In comparison among sorted cells, MMEJ suppression under NHEJ inhibition had minimal effect on the repair patterns, unlike the non-sorted condition. In contrast, treatment with D-I03 in addition to the NHEJi resulted in a significant increase in perfect HDR frequency even in the sorted condition. Interestingly, the augmentation of perfect HDR coincided not with a decrease in deletions as observed in the non-sorted conditions, but with a decrease in imprecise integrations.

To comprehensively analyze the impact of suppressing the two non-HDR pathways on imprecise integration events, we examined the mis-integration patterns under conditions with and without fluorescent sorting. For simplicity, we analyzed three major categories of mis-integration events derived from only single-donor insertion, namely blunt integrations, asymmetric HDR, and imperfect integrations. Among cells in which only the NHEJ was inhibited, the percentage of asymmetric HDR events, where only one side of the donor DNA is precisely integrated, tended to be higher in sorted cells than non-sorted ones. (Fig. 3d). This could be attributed to the scarless insertion of one side of the donor in asymmetric HDR, which may increase the likelihood for functional conservation of the fluorescent protein sequences compared to other patterns of imprecise integrations. In the non-sorted condition, inhibition of the MMEJ or SSA pathway had a marginal impact on the integration pattern (Fig. 3d). In contrast, in the analysis of sorted cells, inhibition of the SSA pathway with D-I03 tended to decrease the percentage of all the three types of imprecise integrations. In particular, asymmetric HDR events, the most common type detected in the sorted cells, exhibited a pronounced tendency for reduction. Collectively, these data suggest that suppression of the SSA pathway increases the proportion of perfect HDR in the fluorescence-positive cells by mainly decreasing the frequency of imprecise integrations, especially asymmetric HDR, in endogenous tagging with fluorescent proteins.

### A flow cytometry-based reporter system further confirmed attenuation of asymmetric HDR upon SSA suppression

To further confirm whether the suppression of the SSA pathway effectively reduces asymmetric HDR, we engineered a flow cytometry-based assay, named the HDR reporter. In this reporter system, when each end of the donor is precisely inserted, cells emit distinct fluorescence wavelengths, which allows for simultaneous evaluation of both perfect and asymmetric HDR events (Fig. 4a). The HDR reporter cassette, containing the N-terminal half of mNG, a spacer sequence, and the C-terminal half of mScarlet, was inserted at the *HNRNPA1* locus in the genome of RPE1 cells. The spacer is flanked by the same unique target sequences for Cas9 placed in the opposite directions, which originate from the mouse *Adenylate kinase 2* gene and exhibit minimal homology to the human genome. The cells were transfected with Cas9-RNP and the donor DNA, which contains the complementary C-terminal half of mNG, the 2A peptides sequence, and the complementary N-terminal half of mScarlet, all flanked by 100 bases of HA sequences. In this system, the seamless repair of the Cas9-induced DSBs in a perfect HDR manner will complete the two split fluorescent protein sequences, resulting in their simultaneous fluorescence emission. Conversely, in cases where indels occurred due to imprecise integrations, mutations would be introduced in the middle of fluorescent protein sequences. Since these mutations would very likely disrupt the functionality of the fluorescent protein, the HDR reporter is highly susceptible to imprecise integrations. Hence, the presence of either mNG or mScarlet fluorescence alone would indicate asymmetric HDR events with precise editing only at the 3’ or 5’ end, respectively.

**Fig. 4:**
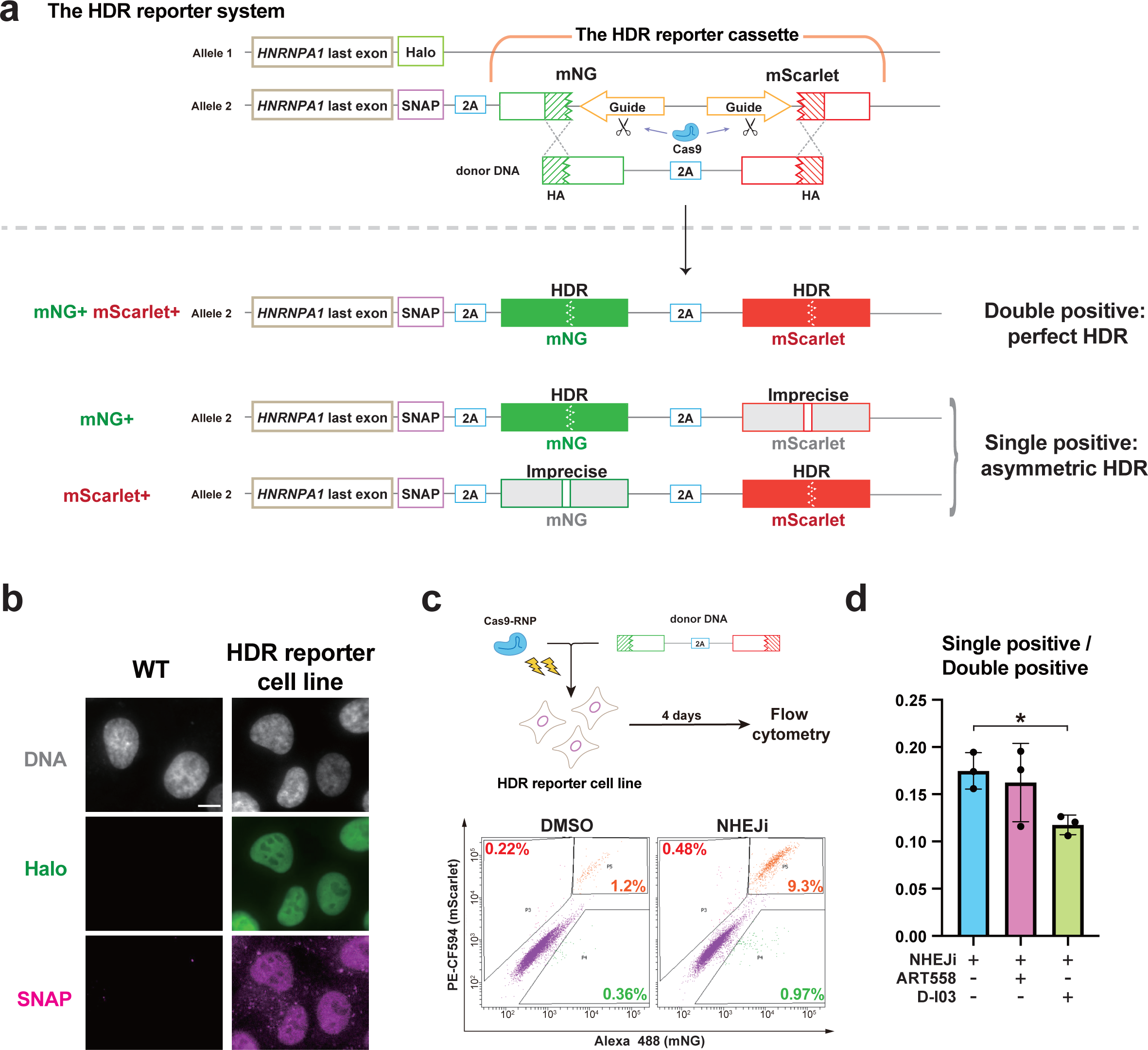
The suppression of SSA pathway reduces asymmetric HDR repair in the HDR reporter assay. **a**, Overview of the HDR reporter system. The HDR reporter cassette is monoallelically inserted at the immediate downstream of the last coding sequence of *HNRNPA1* in RPE1 cells. The seamless, perfect HDR repair of the Cas9-induced DSBs in the reporter cassette completes the two split fluorescent protein sequences, resulting in their simultaneous fluorescence emission (Double positive). The presence of either mNG or mScarlet fluorescence alone (Single positive) shows asymmetric HDR repair with precise editing only at the 3’ or 5’ end, respectively. The sequence of the donor DNA is shown in Supplementary Table 3. **b**, Representative images of the HDR reporter cell line exhibiting both SNAP-tag and HaloTag ligands signals in the localization corresponding to the nuclear protein HNRNPA1. The parental wild type (WT) cells were analyzed as a control. Scale bar: 10 µm. **c**, Time course of the HDR reporter assay and representative plots obtained through flow cytometric analysis. After electroporation, the cells were treated with DMSO or NHEJi for 24 hours. The plots display the percentages of cells exhibiting only mNG signal (green), only mScarlet signal (magenta), or both signals (orange). **d**, The ratio of the single-positive cells to double-positive cells in the HDR reporter assay. Cells were treated with the indicated inhibitors for 24 hours after electroporation. Data from three biological replicates are presented as mean ± S.D. and a two-tailed, unpaired Student’s t-test was used to obtain the P value. *P < 0.05.

To establish the HDR reporter cell line, we utilized the CRISPR/Cpf1 system to insert the reporter cassette into the genome, targeting the C-terminus of HNRNPA1. To use as a readout marker of successful genomic integration, the upstream region of the HDR reporter donor template contains a sequence encoding SNAP-tag, which enables the C-terminus of HNRNPA1 to be endogenously SNAP-tagged (Fig. 4a) Furthermore, in order to exclude mixed signals arising from distinct repair events on the two alleles, we simultaneously introduced a HaloTag donor along with the SNAP-tag-containing HDR reporter donor into cells . Thus, we successfully obtained a cell line with a mono-allelic insertion of the HDR reporter cassette, characterized by exhibiting double positivity for these two ligand-binding tags (Fig. 4b).

Next, we performed knock-in using the donor completing the fluorescent protein sequences in the HDR reporter cell line. Subsequent flow cytometric analysis showed that the major fraction of fluorescence-positive cells exhibited dual fluorescence for both mNG and mScarlet, indicating the occurrence of perfect HDR events (Fig. 4c). Along with that, there was a smaller but considerable portion of single fluorescence-positive cells with only either mNG or mScarlet fluorescence, indicating the presence of asymmetric HDR events. Next, we utilized this reporter system to examine the effects of MMEJ or SSA suppression on the relative occurrence of asymmetric HDR in comparison with perfect HDR. Under NHEJ inhibition, D-I03 treatment decreased the ratio of the single-positive cells to double-positive cells, whereas ART558 had no significant effects (Fig. 4d). This result indicates that SSA suppression specifically reduces asymmetric HDR, consistent with the data of the sequencing analysis using the sorted fluorescence-positive cells (Fig. 3d).

## Discussion

In this study, we comprehensively evaluated the influence of the minor DSB repair pathways on the outcomes of knock-in using CRISPR-mediated endogenous tagging in human diploid RPE1 cells. Quantitative analysis demonstrated that upon hampering NHEJ, the occurrence of repair patterns distinct from perfect HDR is decreased by the inhibition of key factors in the MMEJ and SSA pathways, − POLQ and Rad52, respectively. The suppression of MMEJ and SSA pathways led to a reduction in distinct repair patterns, indicating that they may improve the efficiency of precise knock-in through different mechanisms.

In Cpf1-mediated endogenous tagging of HNRNPA1, the suppression of MMEJ using the POLQ inhibitor reduced overall deletion events and increased perfect HDR frequency upon NHEJ inhibition, consistent with the previous study (Schimmel et al., 2023). Among the deletions, those occurring in the region flanked by two consecutive T bases (five and six in length), with one of them positioned in the 5’ overhang region generated by Cpf1 cleavage, exhibited particularly high susceptibility to POLQ inhibition. This can be attributed to the availability of multiple combinations of microhomology annealing using the two T-rich sequences. Interestingly, Rad52 inhibition demonstrated a more specific reduction in the deletions within the same region. Considering that a homology length of 5 nt at maximum is insufficient to achieve typical DSB repair mediated by the SSA pathway (Ceccaldi et al., 2016; Kelso et al., 2019), it is plausible that the reduced deletions observed upon Rad52 inhibition underlie the suppression of microhomology annealing via the MMEJ pathway. It is possible that the ssDNA-binding of Rad52, which represents the early stages of the SSA pathway, occurs in the 5’ overhang to protect the T-rich sequences that can be utilized by POLQ. Thus, we suggest that Rad52, which plays a major role in the SSA pathway, is also involved in facilitating repairs through the MMEJ pathway by protecting ssDNA, thereby indicating a potential cooperation between these two pathways to promote the repair process. This notion is further supported by the observation that the knock-in efficiency at the *HNRNPA1* locus was synergistically increased by the inhibition of both POLQ and Rad52 (Fig. 2h).

In contrast to *HNRNPA1*, the inhibition of the SSA pathway did not lead to an increase in knock-in efficiency in the other examined genes, which implies deletion events were not decreased at these loci. At the *RAB11A* locus, DNA cleavage was induced by Cas9, which introduces DSBs with blunt ends without ssDNA overhangs (Zaidi et al., 2017). As for the *TOMM20* locus, although Cpf1-mediated cleavage generates DSBs with ssDNA overhangs, there are much fewer microhomologous sequences between the overhangs and its neighboring regions, compared to the *HNRNPA1* locus (Fig. S2). Thus, the correlation between sensitivity to SSA inhibition and the presence of ssDNA overhangs with microhomology to nearby regions supports our hypothesis that Rad52 facilitates introduction of deletions at the lesion via microhomology annealing through ssDNA 5’ overhangs.

Regarding repair patterns associated with insertions of knock-in donors, our observation indicates that the MMEJ pathway has minimal contribution to donor integration (Fig. 3). In the case of random integration of exogenous dsDNA, however, previous studies have shown that MMEJ substantially contributes to genomic insertion of dsDNA (Saito et al., 2017; Zelensky et al., 2017). This difference in contribution may be explained by the presence of HA sequences (90 bp) in the knock-in donors. Notably, POLQ is known to show preference for short microhomology, as it has been reported to be crucial in repair events involving short homologous sequences (6 nt) but not longer ones (≥18 nt) flanking the DSB site (Kelso et al., 2019). Therefore, the presence of long homology regions between the target site and HAs of the donor DNA possibly impedes MMEJ-mediated donor integration.

In contrast to MMEJ, the long-read sequencing analysis showed that imprecise donor integrations in the asymmetric HDR manner were reduced by the SSA pathway suppression, which is further confirmed by the HDR reporter assay. In CRISPR knock-in systems, there is a sufficient length of homology between the HAs of the donor DNA and the target genomic sequence, where annealing of these sequences can be facilitated by the SSA pathway. Through this annealing process, the SSA pathway has been proposed to be potentially utilized for precise donor insertions (Han et al., 2023; Yao et al., 2017; Zhang et al., 2017), although to our knowledge, there was no prior experimental evidence to support this hypothesis. The data obtained in this study suggest that SSA can directly mediate donor insertion into CRISPR/Cas-mediated DSBs, but particularly in an asymmetric HDR manner. In other words, the SSA pathway is likely to facilitate precise donor insertion at only one of its end, while exhibiting limited contribution to the occurrence of perfect HDR events, where both of the donor ends are precisely inserted. Rather, our data suggest that SSA inhibition has favorable effects on augmenting perfect HDR events under specific conditions (Fig. 3c, d).

When practically performing CRISPR/Cas-mediated knock-in, how the induced DSBs are repaired can be influenced by the genomic environment of the specific target loci and the experimental system being utilized. Indeed, our study provides evidence that the distribution of repair outcomes, as well as the impact of inhibiting DSB repair pathways, can vary depending on the target gene locus and the type of Cas nuclease used. Moreover, in the case of endogenous tagging with fluorescent proteins, the presence or absence of fluorescence sorting procedure affected the ultimate distribution of repair patterns and thus outcomes of repair pathway inhibition. Considering these contextual factors, we propose that the efficiency of precise knock-in can be effectively enhanced by appropriately employing tailored combinations of DSB repair pathway inhibition.

## Acknowledgments

We thank Miho Kiyooka and Wei Chen at National Institute of Genetics for supporting PacBio sequencing, Dr. Yusuke Kishi at Institute for Quantitative Biosciences at the University of Tokyo for supporting quality control of PacBio library preparation, and the Kitagawa lab members for technical supports and helpful discussions.

## Author contributions

S.H. conceived and designed the study. C.T. designed and performed most of the experiments. A.M. and M.G. optimized the genome editing conditions. A.M. and C.T. designed the HDR reporter system. S.O. validated the completion of the HDR reporter cell clone. C.T., A.M. and K.K.I. analyzed the PacBio data with *knock-knock*. M.F., S.Y. and T.C. provided suggestions. A.T. performed PacBio sequencing. C.T., S.H., A.M. and D.K. analyzed the data. C.T., A.M., S.H. and D.K. wrote the manuscript. All authors contributed to discussions and manuscript preparation.

## Material and methods

### Cell culture

RPE1 cells obtained from the American Type Culture Collection (ATCC) were maintained in Dulbecco’s Modified Eagle Medium/Nutrient Mixture F-12 (DMEM/F-12, Nacalai Tesque) with 10% FBS, 100 U/mL penicillin, and 100 µg/mL streptomycin, respectively. Cells were cultured at 37°C in a humidified 5% CO2 incubator.

### Chemicals

Alt-R HDR enhancer V2 (referred to as NHEJi) from Integrated DNA Technologies (IDT) was used as NHEJi at 1 μM and stored at -20°C according to the manufacturer’s guidelines. ART558 (MedChemExpress) and D-I03 (MedChemExpress) were dissolved as 10 mM stock in DMSO, stored at -80°C, and used with a final concentration of 10 μM in medium.

### DNA donor and guide RNA preparation

DNA donors and guide RNAs (sgRNA for Cas9 and crRNA for Cpf1) were prepared according to previously published methods (Komori et al., 2023; Mabuchi et al., 2023). Donor DNAs for C-terminal tagging, containing the 5xGA linker-mNG/mScarlet sequence, and for N-terminal tagging, containing the mNG/mScarlet-5xGA linker sequence, were amplified by PCR from plasmids encoding respective sequences. Two primers were used, each containing a 90-base left or right homology arm (HA) sequence. We used Q5 High-Fidelity 2X Master Mix (New England Biolabs) for PCR. DpnI and exonuclease I were used for digestion of residual template plasmids and primers, respectively. The DNA donors were then column-purified using the NucleoSpin Gel and PCR Clean-up kit (Macherey-Nagel) and stored at -20°C or directly used for electroporation. As for mNG tagging of CLTA, donor DNA was prepared with additional 2nd-round PCR using a gel-purified 1st PCR product as a template to minimize non-specific products. All primer sequences used in this study are listed in Supplementary Table 1. A guide RNA (sgRNA for Cas9 and crRNA for Cpf1) was transcribed *in vitro* from PCR-generated DNA templates. A template DNA containing T7 promoter and sgRNA sequence was amplified by PCR. After treatment of DNase I (Takara Bio), the synthesized guide RNA was purified using the RNA Clean & Concentrator Kit (Zymo Research). All target site sequences of guide RNAs used in this study are listed in Supplementary Table 2.

### Gene knock-in using the CRISPR/Cpf1 and CRISPR/Cas9 system

Endogenous gene tagging using the CRISPR-Cpf1 system was performed with the electroporation of Cpf1-RNP and DNA donors using the Neon Transfection System (Thermo Fisher Scientific) according to the manufacturer’s protocol. A.s. Cas12a Ultra (1 µM) (Cpf1) from Integrated DNA Technologies (IDT) and crRNA (1 µM) were pre-incubated in resuspension buffer R (Thermo Fisher Scientific) at room temperature and mixed with cells (0.125 ×10^5^ /µL), Cpf1 electroporation enhancer (1.8 µM, IDT), and single or dual DNA donors (33 nM in total, i.e., 16.5 nM for each DNA donor in dual color tagging). Electroporation was conducted using a 10 µL Neon tip at a voltage of 1300 V with two 20 ms pulses. The transfected cells were seeded into a 24-well plate with medium containing inhibitors. 24 h after electroporation, the inhibitors were removed by replacing the culture medium three times.

CRISPR/Cas9-mediated knock-in was performed similarly to the Cpf1-RNP condition described above, with a modification in the electroporation solution. Briefly, HiFi Cas9 protein (1.55 µM, IDT) and sgRNA (1.84 µM) were pre-incubated in buffer R and mixed with cells, Cas9 electroporation enhancer (1.8 µM, IDT), and DNA donors. Electroporation was conducted at a voltage of 1300 V with two 20 ms pulses.

### Fluorescent protein imaging in CRISPR-mediated mNG tagging of HNRNPA1 and RAB11A

5 days after electroporation, cells cultured on coverslips (Matsunami) were fixed with 4% PFA at room temperature for 15 min. After PBS washing, fixed cells were subjected to permeabilization with PBS containing 0.05% Triton X-100 for 5 min at room temperature. The coverslips were washed with PBS and mounted onto glass slides (Matsunami) using ProLong Gold Antifade Mountant with DAPI (Invitrogen), with the cell side down. The representative images were acquired using Axio Imager.M2 microscope (Carl Zeiss) equipped with a 63× lens objective.

### Quantification of knock-in efficiency by flow cytometry

Flow cytometric analysis was conducted 4 days after electroporation. Cells were harvested with trypsin/EDTA solution and suspended in PBS. The cell suspensions were analyzed using BD FACS Aria III (BD Biosciences), equipped with 355/405/488/561/633 nm lasers to detect cells with mNG or mScarlet signal. Data were collected from more than 5000 gated events.

### siRNA-mediated gene knockdown

The following siRNAs were used: Silencer Select siRNA (Life Technologies) against POLQ no.1 (s21059), POLQ no.2 (s21060), Rad52 no. 1 (s11546), Rad52 no. 2 (s11747), and negative control (4390843).

Lipofectamine RNAiMAX (Thermo Fisher Scientific) was used with a final concentration of 20 nM total siRNA (Two siRNAs against the same gene were mixed to a final concentration of 10 nM, respectively) for siRNA transfection according to manufacturer’s protocol. Transfected cells were cultured for 24 h before electroporation.

### Amplicon sequencing and analysis by *knock-knock* and SIQ

#### Genomic DNA preparation

For amplicon sequencing analysis conducted without the flow cytometric sorting, after electroporation of Cas-RNP targeting *HNRNPA1*, *TOMM20*, *RAB11A* or *CLTA* loci and a donor DNA, the cells were cultured in media containing inhibitors for 24 h, followed by an additional 3 days culture in fresh media. Genomic DNA was then extracted using NucleoSpin DNA RapidLyse kit (Macherey-Nagel).

For amplicon sequencing analysis performed for sorted fluorescence-positive cells, following electroporation of Cpf1-RNP targeting *HNRNPA1* locus and the donor DNA, the cells were cultured in media with inhibitors for 24 hours. After subsequent incubation in fresh media for 3 days, 10,000 mNG positive cells were collected using FACS Aria III, followed by genomic DNA extraction using NucleoSpin Tissue XS kit (Macherey-Nagel).

#### Amplicon sequencing

Amplicon libraries were generated using a 2-step PCR and adapter ligation protocol based on the instructions by Pacific Biosciences (Part Number 101-791-800 Version 02, April 2020) with slight modifications. In the first round of PCR, a region flanking the target site of mNG insertion was amplified from the extracted genomic DNA using KOD One Master Mix (TOYOBO) and primers with universal sequences that provide an annealing site for a barcoded primer. These universal sequences served as annealing sites for barcoded primers. The amplified DNA was purified using AMPure XP beads (Beckman Coulter). Subsequently, the purified DNA underwent a second round of PCR using primers from the Barcoded Universal F/R Primers Plate-96v2 (Pacific Biosciences), followed by purification using AMPure PB beads (Pacific Biosciences). The barcoded amplicons were analyzed using TapeStation (Agilent Technologies) and Qubit Fluorometer (Thermo Fisher Scientific). Finally, all the amplicons were pooled together in equimolar amounts as a single sample. A pooled sequencing libraries were prepared using the SMRTbell Express Template Prep Kit 2.0 (Pacific Biosciences, CA, USA). Three Sequel II SMRT cells were run on the PacBio Sequel II system with Binding Kit 3.1 or 3.2/Sequencing Kit 2.0 (Pacific Biosciences, CA, USA) and 30 hour movies. The consensus (HiFi) reads were generated from the raw full-pass subreads using the DeepConsensus v1.1.0 or v1.2 program (Baid et al., 2022), and then were demultiplexed by sample barcodes using the SMRT Link v11.0.0.146107 software (Pacific Biosciences, CA, USA). A total of 5,181,459 barcoded Reads with read quality score >= 40 were selected. The information about the sequencing results were shown in Supplementary Table 4.

#### Analysis of knock-in outcomes by knock-knock

After trimming the universal sequences from the reads, the trimmed reads were subjected to analysis using *knock-knock*, a computational pipeline developed by Canaj et al. (2019), to examine the knock-in outcomes. We used a software package of *knock-knock* available at https://github.com/jeffhussmann/knock-knock. Based on the detailed classification provided by *knock-knock,* repair patterns that were classified as "5’ blunt, 3’ blunt" were grouped as "blunt". Repair patterns classified as "5’ HDR, 3’ blunt", "5’ blunt, 3’ HDR", "5’ HDR, 3’ imperfect", and "5’ imperfect, 3’ HDR" were categorized as "asymmetric HDR". Repair patterns classified as "5’ blunt, 3’ imperfect", "5’ imperfect, 3’ blunt", and "5’ imperfect, 3’ imperfect" were assigned to the "imperfect" category in Figure 1 and 3.

#### Analysis of deletion patterns by SIQ

Reads classified to deletions of less than 50 nt by *knock-knock* were further analyzed by Sequence Interrogation and Quantification (SIQ) program (van Schendel et al., 2022). SIQ provides a comprehensive analysis of a mutation profile at the target site that can be visualized using SIQPlotteR. We used a software package available at https://github.com/RobinVanSchendel/SIQ.

### Establishment of the HDR reporter

#### Preparation of donor DNA for knock-in of the HDR reporter cassette to HNRNPA1

Donor DNA for knock-in of the HDR reporter cassette was prepared by PCR using a plasmid that had been cloned to include SNAP-tag, 2A peptide, and the HDR reporter cassette in this order flanked by HAs for knock-in at the *HNRNPA1* locus. The HaloTag donor was PCR-amplified from a plasmid encoding the 5xGA-HaloTag sequence using two primers containing left and right HAs for knock-in at the *HNRNPA1* locus. The sequence of the donor DNA containing SNAP-tag, 2A peptide, and the HDR reporter cassette are listed in Supplementary Table 3.

#### The design of the guide sequence for Cas9

The guide sequence candidates were obtained by searching for guide sequences targeting some mouse genes using the CRISPR-Cas9 guide RNA design checker provided by IDT. Subsequently, the frequency of off-target sequences in the human genome was assessed for each guide sequence using the CRISPR-Cas9 guide RNA design checker, and the guide sequence targeting mouse *Adenylate kinase 2* with the minimal off-targets was chosen (Supplementary Table 2).

#### Isolation of single cells

A few weeks after electroporation of Cpf1-RNP targeting the *HNRNPA1* locus and two DNA donors (The HDR reporter cassette donor and HaloTag donor), cells were labeled with HaloTag Oregon Green (Promega) according to the manufacturer’s protocol. Halo-positive single cells were isolated into a 96-well plate using BD FACS Aria III Cell Sorter.

After 17 days culture, the cell clones were seeded into two 96-well plates, one for expansion and the other for visualization. On the following day, cells in the latter plate were labeled with HaloTag Oregon Green and SNAP-Cell 647-SiR (New England Biolabs) simultaneously according to the manufacturers’ protocols. SNAP/Halo double-positive clones were selected after observation using CQ1 Benchtop High-Content Analysis System (Yokogawa Electric Corp). After cell expansion, the SNAP/Halo-positive clone cultured on coverslips (Matsunami Glass) were treated with HaloTag Oregon Green and SNAP-Cell TMR-Star (New England Biolabs) according to manufacturers’ protocols and fixed with 4% PFA in PBS at room temperature for 15 min. The coverslips with fixed cells were washed with PBS and mounted onto glass slides using ProLong Gold Antifade Mountant with DAPI, with the cell side down. The representative images of the HDR reporter cell line were acquired using Axio Imager.M2 microscope equipped with a 63× lens objective.

### Statistical analysis

Statistical comparison between the data from different groups was performed in PRISM v.9 software (GraphPad) using either a Tukey–Kramer test or a two-tailed, unpaired Student’s t test as indicated in the figure legends. P values <0.05 were considered statistically significant. All data shown are mean ± S.D. Sample sizes are indicated in the figure legends.

