## Supplementary Tables for "Comparable analysis of multiple DNA double-strand break repair pathways in CRISPR-mediated endogenous tagging"

#### Supplementary Table 1: Primer sequences for PCR

##### For guide RNA assembly

| Name | Sequence |
|---|---|
| Cpf1_crRNA_Fw | TTCTAATACGACTCACTATAGTAATTTCTACTCTTGTAGAT |
| HNRNPA1_crRNA_Rv | AAGGTGCTTACTTACCTAATATCTACAAGAGTAGAAATTAC |
| TOMM20_crRNA_Rv | CTGAAGATGATGTGGAATGAATCTACAAGAGTAGAAATTAC |
| Univ_sgRNA_Fw | TTCTAATACGACTCACTATAG |
| Univ_sgRNA_Rv | AAAAGCACCGACTCGGTG |
| crRNA_tracrRNA | GTTTTAGAGCTAGAAATAGCAAGTTAAAATAAGGCTAGTCCGTTATCAACTTGAAAAAGTGGCACCGAGTCGGTGCTTTT |
| RAB11A_sgRNA_Fw | TTCTAATACGACTCACTATAGGTAGTCGTACTCGTCGTCG |
| RAB11A_sgRNA_ Rv | TTCTAGCTCTAAAACCGACGACGAGTACGACTACC |
| CLTA_sgRNA_Fw | TTCTAATACGACTCACTATAGAACGGATCCAGCTCAGCCA |
| CLTA_sgRNA_Rv | TTCTAGCTCTAAAACTGGCTGAGCTGGATCCGTTC |

##### For HDR donor preparation

| Name | Sequence |
|---|---|
| HNRNPA1-mNG/mScarlet_Fw | CACTTTGAAACTTTAAAAGAAAAATTGTACTTTTCAGGTGGCTATGGCGGTTCCAGCAGCAGCAGTAGCTATGGCAGTGGCAGAAGATTTGGAGCTGGTGCAGGTGCAG |
| HNRNPA1-mNG/mScarlet_Rv | ACTGCAATTATAATGTTAACTATGTTGCACTGCTCAGCTACATTAGGGTTATTGGGTTCATCAGCAATTTAAAAAATTATGTCAACACACAAAAAGGTGCTTACTTACCTAACTACTTGTACAGCTCGTCCATGC |
| TOMM20-mNG/mScarlet_Fw | TATTTTGAAGTTAGAATCCTAATTAAATGCTTATGACACTTTAAAAAATTATTTTTTTTTCTTTCAGAGAATTGTAAGTGCTCAGAGCTTGGCTGAAGATGATGTGGAAGGAGCTGGTGCAGGTGCAG |
| TOMM20-mNG/mScarlet_Rv | ATATTTGCCCTTATTCCCCCAGAGCTGCTCAACTACCAAGAATTTTTAAAATATTTTTAACTGAGATTTTATTATGTTGACATTTGTTTCCTACTTGTACAGCTCGTCCATGC |
| mNG-RAB11A_Fw | GGCGCTCGGGTTACCCCTGCAGCGACGCCCCCTGGTCCCACAGATACCACTGCTGCTCCCGCCCTTTCGCTCCTCGGCCGCGCAATGGGCATGGTGAGCAAGGGCGAG |
| mScarlet-RAB11A_Fw | GGCGCTCGGGTTACCCCTGCAGCGACGCCCCCTGGTCCCACAGATACCACTGCTGCTCCCGCCCTTTCGCTCCTCGGCCGCGCAATGGGCATGGTGAGCAAGGGCGAGGC |
| mNG/mScarlet-RAB11A_Rv | GGGAGTGGCCCGGGTCCCCGAACGAGGACTGTGTAGAGTGCGAGAGCCCATGGCCTCACCTTTAAAGAGGTAGTCGTACTCGTCGTCGCGTGCACCAGCTCCTGCACC |
| mNG-CLTA_Fw | ACAGCGGTGGCTGCCGGGCGTGGTGTCGGTGGGTCGGTTGGTTTTTGTCTCACCGTTGGTGTCCGTGCCGTTCAGTTGCCCGCCATGGCTATGGTGAGCAAGGGCGAG |

| | |
|---|---|
| mScarlet-CLTA_Fw | ACAGCGGTGGCTGCCGGGCGTGGTGTCGGTGGGTCGGTTGGTTTTTGTCTCACCGTTGGTGTCCGTGCCGTTCAGTTGCCCGCCATGGCTATGGTGAGCAAGGGCGAGGC |
| mNG/mScarlet-CLTA_Rv | AGCCGGGTCTTCTTCGCCGGCGCCGGCCACTCCGTTCCCCAGCGCGGACCGCCAGGGGCGCCGGCAGGGGCGCCGAACGGATCCAGCTCTGCACCAGCTCCTGCACC |
| mNG-CLTA_2nd_Fw | ACAGCGGTGGCTGCCGGG |
| mNG-CLTA_2nd_Rv | AGCCGGGTCTTCTTCGCCGG |
| The HDR reporter donor_Fw | CCGCCATGGTAGATGGCTCC |
| The HDR reporter donor_Rv | CCGCTCGGTGGACGCTTC |
| HDR reporter cassette knock-in_Fw | CTTGTAGTAGGGCCATTTTTAAATGGCCAGACACTTGAATTTAACTTTTATTATCCCAAATATGAAAACATTACTGTTGGCACTTTGAAACTTTAAAAGAAAAATTGTACTTTTCAGGTGGCTATGGCGGTTCCAGCAG |
| HDR reporter cassette knock-in_Rv | AGACTCAAGGCTACAATCCAATATCAAGTTTGTTTCCCACAAATTTTGCTGATCTGAATATTAACTTTATATCCACAATTACTGCAATTATAATGTTAACTATGTTGCACTGCTCAGCTACATTAGGGTTATTGGGTTCATCAGC |
| HNRNPA1-HaloTag_Fw | CACTTTGAAACTTTAAAAGAAAAATTGTACTTTTCAGGTGGCTATGGCGGTTCCAGCAGCAGCAGTAGCTATGGCAGTGGCAGAAGATTTGGAGCTGGTGCAGGTGCAG |
| HNRNPA1-HaloTag_Rv | ACTGCAATTATAATGTTAACTATGTTGCACTGCTCAGCTACATTAGGGTTATTGGGTTCATCAGCAATTTAAAAAATTATGTCAACACACAAAAAGTTGCTTACTTACCTAACTAGCCGGAAATCTCGAGCG |

#### For long-read amplicon sequencing

| Name | Sequence |
|---|---|
| SMRT_1st_Fw_HNRNPA1 | [AmC6]GCAGTCGAACATGTAGCTGACTCAGGTCACCAGGCCTTCAGCCGTTACAC |
| SMRT_1st_Rv_HNRNPA1 | [AmC6]TGGATCACTTGTGCAAGCATCACATCGTAGCCCAACCAGAACCCAGTCAAACT |
| SMRT_1st_Fw_TOMM20 | [AmC6]GCAGTCGAACATGTAGCTGACTCAGGTCACTGATCTGCCTCCTTTGTTAACTTG |
| SMRT_1st_Rv_TOMM20 | [AmC6]TGGATCACTTGTGCAAGCATCACATCGTAGCTAGCGAAGCTCACAAGGCT |
| SMRT_1st_Fw_RAB11A | [AmC6]GCAGTCGAACATGTAGCTGACTCAGGTCACGCAGTGAAGAAGCTCATTAAGACAAC |
| SMRT_1st_Rv_RAB11A | [AmC6]TGGATCACTTGTGCAAGCATCACATCGTAGGAAGGTAGAGAGAGTTGCCAAATGG |
| SMRT_1st_Fw_CLTA | [AmC6]GCAGTCGAACATGTAGCTGACTCAGGTCACAGCCATGTAGCTATTAACATCTCCCTG |
| SMRT_1st_Rv_CLTA | [AmC6]TGGATCACTTGTGCAAGCATCACATCGTAGCCAACACTCTGTACACCTTAAGTGC |

#### Supplementary Table 2: Target site sequences of guide RNA

| Target gene | Cas nuclease | Target sequence |
|---|---|---|
| *HNRNPA1* (Human) | Cpf1 | ATTAGGTAAGTAAGCACCTT |
| *TOMM20* (Human) | Cpf1 | TCATTCCACATCATCTTCAG |
| *RAB11A* (Human) | Cas9 | GGTAGTCGTACTCGTCGTCG |
| *CLTA* (Human) | Cas9 | GAACGGATCCAGCTCAGCCA |
| *Adenylate kinase 2* (Mouse) | Cas9 | CGGTTCGGAAGCCAACACGT |

#### Supplementary Table 3: The sequences of the donor DNAs related to the HDR reporter system

| Description | Sequence |
|---|---|
| The donor DNA in the HDR reporter assay | CCGCCATGGTAGATGGCTCCGGATACCAAGTCCATCGCACAATGCAGTTTGA<br>AGATGGTGCCTCCCTTACTGTTAACTACCGCTACACCTACGAGGGAAGCCAC<br>ATCAAAGGAGAGGCCCAGGTGAAGGGGACTGGTTTCCCTGCTGACGGTCCT<br>GTGATGACCAACTCGCTGACCGCTGCGGACTGGTGCAGGTCGAAGAAGACT<br>TACCCCAACGACAAAACCATCATCAGTACCTTTAAGTGGAGTTACACCACTGG<br>AAATGGCAAGCGCTACCGGAGCACTGCGCGGACCACCTACACCTTTGCCAA<br>GCCAATGGCGGCTAACTATCTGAAGAACCAGCCGATGTACGTGTTCCGTAAG<br>ACGGAGCTCAAGCACTCCAAGACCGAGCTCAACTTCAAGGAGTGGCAAAAG<br>GCCTTTACCGATGTGATGGGCATGGACGAGCTGTACAAGGGCAGCGGCGCC<br>ACAAACTTCTCTCTGCTAAAGCAAGCAGGTGATGTTGAAGAAAACCCCGGGC<br>CTATGGTGAGCAAGGGCGAGGCAGTGATCAAGGAGTTCATGCGGTTCAAGG<br>TGCACATGGAGGGCTCCATGAACGGCCACGAGTTCGAGATCGAGGGCGAG<br>GGCGAGGGCCGCCCCTACGAGGGCACCCAGACCGCCAAGCTGAAGGTGAC<br>CAAGGGTGGCCCCCTGCCCTTCTCCTGGGACATCCTGTCCCCTCAGTTCAT<br>GTACGGCTCCAGGGCCTTCATCAAGCACCCCGCCGACATCCCCGACTACTAT<br>AAGCAGTCCTTCCCCGAGGGCTTCAAGTGGGAGCGCGTGATGAACTTCGAG<br>GACGGCGGCGCCGTGACCGTGACCCAGGACACCTCCCTGGAGGACGGCA<br>CCCTGATCTACAAGGTGAAGCTCCGCGGCACCAACTTCCCTCCTGACGGCC<br>CCGTAATGCAGAAGAAGACAATGGGCTGGGAAGCGTCCACCGAGCGG |
| The donor DNA used to insert the HDR reporter cassette into the *HNRNPA1* locus | CTTGTAGTAGGGCCATTTTTAAATGGCCAGACACTTGAATTTAACTTTTATTATC<br>CCAAATATGAAAACATTACTGTTGGCACTTTGAAACTTTAAAAGAAAAATTGTA<br>CTTTTCAGGTGGCTATGGCGGTTCCAGCAGCAGCAGTAGCTATGGCAGTGG<br>CAGAAGATTTTCAGGTGGAGGCGGTTCAGGCGGAGGTGGCTCTGGCGGTG<br>GCGGATCCATGGACAAGGATTGTGAAATGAAACGCACCACACTGGACAGCC<br>CTTTGGGGAAGCTGGAGCTGTCTGGTTGTGAGCAGGGTCTGCACGAAATAA<br>AGCTCCTGGGCAAGGGGACGTCTGCAGCTGATGCCGTGGAGGTCCCAGCC<br>CCCGCTGCGGTTCTCGGAGGTCCGGAGCCCCTGATGCAATGCACAGCCTG<br>GCTGAATGCCTATTTCCACCAGCCCGAGGCTATCGAAGAGTTCCCCGTGCCG<br>GCTCTTCACCATCCCGTTTTCCAGCAAGAGTCGTTCACCAGACAGGTGTTAT<br>GGAAGCTGCTGAAGGTTGTGAAATTCGGAGAAGTGATTTCTTACCAGCAATT<br>AGCAGCCCTGGCAGGCAACCCCGCAGCCACGGCAGCAGTGGGAGGAGCA<br>ATGAGAGGCAATCCTGTCCCTATCCTGATCCCGTGCCACAGAGTGGTCTGCA<br>GCAGCGGAGCCGTGGGCGGTTACGAGGGTGGACTGGCCGTGAAGGAATGG<br>CTTCTGGCCCATGAAGGCCATCGGTTGGGGAAGCCAGGCTTGGGAGGCTC<br>CGGCGAGGGCAGGGGAAGTCTTCTAACATGCGGGGACGTGGAGGAAAATC<br>CCGGCCCAATGGTGAGCAAGGGCGAGGAGGATAACATGGCCTCTCTCCCAG<br>CGACACATGAGTTACACATCTTTGGCTCCATCAACGGTGTGGACTTTGACAT<br>GGTGGGTCAGGGCACCGGCAATCCAAATGATGGTTATGAGGAGTTAAACCTG<br>AAGTCCACCAAGGGTGACCTCCAGTTCTCCCCCTGGATTCTGGTCCCTCATA<br>TCGGGTATGGCTTCCATCAGTACCTGCCCTACCCTGACGGGATGTCGCCTTT<br>CCAGGCCGCCATGGTAGATGGCTCCGGATACCAAGTCCATCGCACAATGCA<br>GTTTGAAGATGGTGCCTCCCTTACTGTTAACTACCGCTACACCTACGAGGGA<br>AGCCTACGTGTTGGCTTCCGAACCGAAGATATCTAGGGATAACAGGGTAATC<br>CCGGGGGCGGTTCGGAAGCCAACACGTAGGCACCCTGATCTACAAGGTGAA<br>GCTCCGCGGCACCAACTTCCCTCCTGACGGCCCCGTAATGCAGAAGAAGAC |

| | |
|---|---|
| | AATGGGCTGGGAAGCGTCCACCGAGCGGTTGTACCCCGAGGACGGCGTGC<br>TGAAGGGCGACATTAAGATGGCCCTGCGCCTGAAGGACGGAGGCCGCTACC<br>TGGCGGACTTCAAGACCACCTACAAGGCCAAGAAGCCCGTGCAGATGCCCG<br>GCGCCTACAACGTCGACCGCAAGTTGGACATCACCTCCCACAACGAGGACT<br>ACACCGTGGTGGAACAGTACGAACGCTCCGAGGGCCGCCACTCCACCGGC<br>GGCATGGACGAGCTGTACAAGTAGTTAGGTAAGTAAGCACCTTTTTGTGTGTT<br>GACATAATTTTTTAAATTGCTGATGAACCCAATAACCCTAATGTAGCTGAGCAG<br>TGCAACATAGTTAACATTATAATTGCAGTAATTGTGGATATAAAGTTAATATTCA<br>GATCAGCAAAATTTGTGGGAAACAAACTTGATATTGGATTGTAGCCTTGAGTC<br>T |

### Supplementary Table 4: The information about the sequencing results

| Run Name | Reagents | Sample Name | Barcode | No. of HiFi reads (>=QV40) | Total Bases (bp) | DeepConsensus | Figure |
|---|---|---|---|---|---|---|---|
| amplicon_230510_spk3bc_20230517_run01 | Binding Kit3.1/Sequencing Kit2.0 | HNRNPA1-1 | bc1072--bc1072 | 209,322 | 549,828,648 | v1.2 | 1 |
| | | HNRNPA1-2 | bc1073--bc1073 | 122,496 | 324,759,844 | | 1 |
| | | HNRNPA1-3 | bc1074--bc1074 | 171,461 | 452,522,572 | | 1 |
| | | TOMM20-1 | bc1075--bc1075 | 131,724 | 147,605,594 | | 1 |
| | | TOMM20-2 | bc1076--bc1076 | 211,434 | 241,442,654 | | 1 |
| | | TOMM20-3 | bc1077--bc1077 | 184,362 | 207,379,789 | | 1 |
| | | RAB11A-1 | bc1084--bc1084 | 203,958 | 311,548,040 | | 1 |
| | | RAB11A-2 | bc1079--bc1079 | 175,255 | 190,005,966 | | 1 |
| | | RAB11A-3 | bc1080--bc1080 | 124,304 | 311,548,040 | | 1 |
| | | CLTA-1 | bc1081--bc1081 | 146,635 | 279,196,586 | | 1 |
| | | CLTA-2 | bc1082--bc1082 | 205,758 | 388,830,301 | | 1 |
| | | CLTA-3 | bc1083--bc1083 | 165,855 | 315,453,488 | | 1 |
| amplicon_230602_spk3bc_20230606_run01 | Binding Kit3.2/Sequencing Kit2.0 | HNRNPA1_NHEJi-1 | bc1046--bc1046 | 157,645 | 481,181,496 | v1.2 | 3 (non-sorted) |
| | | HNRNPA1_NHEJi-2 | bc1047--bc1047 | 145,841 | 453,348,742 | | 3 (non-sorted) |
| | | HNRNPA1_NHEJi-3 | bc1048--bc1048 | 151,343 | 454,731,939 | | 3 (non-sorted) |
| | | HNRNPA1_ART558-1 | bc1049--bc1049 | 230,164 | 723,083,080 | | 3 (non-sorted) |
| | | HNRNPA1_ART558-2 | bc1050--bc1050 | 193,847 | 605,100,475 | | 3 (non-sorted) |
| | | HNRNPA1_ART558-3 | bc1064--bc1064 | 174,940 | 554,503,991 | | 3 (non-sorted) |
| | | HNRNPA1_D-I03-1 | bc1052--bc1052 | 165,286 | 517,254,021 | | 3 (non-sorted) |
| | | HNRNPA1_D-I03-2 | bc1053--bc1053 | 166,622 | 518,457,349 | | 3 (non-sorted) |
| | | HNRNPA1_D-I03-3 | bc1054--bc1054 | 115,790 | 360,358,701 | | 3 (non-sorted) |
| amplicon_230327_spk3bc_20230328_run01 | Binding Kit3.2/Sequencing Kit2.0 | HNRNPA1_NHEJi-1 | bc1002--bc1002 | 183,702 | 604,629,764 | v1.1.0 | 3 (sorted) |
| | | HNRNPA1_NHEJi-2 | bc1007--bc1007 | 157,317 | 519,350,435 | | 3 (sorted) |
| | | HNRNPA1_NHEJi-3 | bc1012--bc1012 | 226,842 | 746,476,738 | | 3 (sorted) |
| | | HNRNPA1_ART558-1 | bc1003--bc1003 | 196,337 | 651,542,282 | | 3 (sorted) |
| | | HNRNPA1_ART558-2 | bc1008--bc1008 | 163,395 | 545,035,511 | | 3 (sorted) |
| | | HNRNPA1_ART558-3 | bc1013--bc1013 | 167,824 | 559,259,524 | | 3 (sorted) |
| | | HNRNPA1_D-I03-1 | bc1004--bc1004 | 200,985 | 662,219,143 | | 3 (sorted) |
| | | HNRNPA1_D-I03-2 | bc1009--bc1009 | 183,749 | 604,523,605 | | 3 (sorted) |
| | | HNRNPA1_D-I03-3 | bc1014--bc1014 | 147,266 | 483,730,873 | | 3 (sorted) |
| | | | Total | 5,181,459 | 13,764,909,191 | | |
